# Molecular characterization of extended-spectrum beta-lactamase-producing extra-intestinal pathogenic *Escherichia coli* isolated in a university teaching hospital Dakar-Senegal

**DOI:** 10.1101/2022.07.20.500880

**Authors:** Komla Mawunyo Dossouvi, Bissoume Sambe Ba, Gora Lo, Abdoulaye Cissé, Awa Ba-Diallo, Issa Ndiaye, Assane Dieng, Serigne Mbaye Lo Ndiaye, Cheikh Fall, Alioune Tine, Farba Karam, Habsa Diagne-Samb, Safietou Ngom-Cisse, Halimatou Diop-Ndiaye, Coumba Toure-Kane, Aïssatou Gaye-Diallo, Souleymane Mboup, Cheikh Saad Bouh Boye, Yakhya Dièye, Abdoulaye Seck, Makhtar Camara

## Abstract

Extra-intestinal pathogenic *Escherichia coli* (ExPEC), a predominant Gram-negative bacterial pathogen, express a wide range of virulence factors and is responsible of several diseases including urinary tract infections (UTI), nosocomial pneumonia, bacteremia, and neonatal meningitis. ExPEC isolates are often multidrug resistant (MDR) and clones producing extended-spectrum beta-lactamases (ESBL) are increasingly reported all over the world.

Seventy-eight clinical ExPEC strains were selected for this study. The majority was from UTIs (n=51), while the rest (n=27) was from pus, sputum, bronchial fluid and vaginal samples (non-uropathogenic ExPEC). Interestingly, 49 out of the 78 ExPEC isolates where considered as community-acquired (CA) and 29 hospital-acquired (HA) bacteria. Antibiotic susceptibility testing was performed using the Kirby-Bauer disc diffusion method. Standard polymerase chain reaction (PCR) was used to screen major ESBL genes (*bla*_CTX-M_, *bla*_OXA-1_, *bla*_TEM_, *bla*_SHV_) and _blaCTX-M_ variants (*bla*_CTX-M-1_, *bla*_CTX-M-9_, *bla*_CTX-M-15_, *bla*_CTX-M-25_).

All the ExPEC strains were resistant to ampicillin, ticarcillin, amoxicillin/clavulanic acid combination, cefalotin, cefotaxime, ceftazidime, cefepime and aztreonam, but showed a high susceptibity to fosfomycin (98.7%, *n* = 77), ertapenem (96.2%, *n* = 75), and imipenem (100%). Moreover, isolates harbored at least one ESBL gene, including *bla*_CTX-M_ (98.7%), *bla*_OXA-1_ (78.2%), *bla*_TEM_ (44.9%) and *bla*_SHV_ (3.8%). The CTX-M variants were also found with the predominance of *bla*_CTX-M-1_ (90.9%) and *bla*_CTX-M-15_ (90.9%) followed by *bla*_CTX-M-9_ (11.7%), while *bla*_CTX-M-25_ was not detected.

Despite the resistance to most of the tested antibiotics, ExPEC isolates showed fortunately a good susceptibility to fosfomycin and carbapenems. *bla*_CTX-M_ (*bla*_CTX-M1_, *bla*_CTX-M15_) and *bla*_OXA-1_ seem to be *E. coli* major ESBL genes circulating in Senegal. No significant difference was noted when comparing prevalence of ESBL genes detected from CA and HA strains, and from UPEC and non-uropathogenic ExPEC. The high level of resistance to antimicrobials observed stresses the need of establishing an epidemiological surveillance of antimicrobial resistance in both community and hospital settings.

## Introduction

*Escherichia coli* (*E. coli*), a common bacteria found in various parts of the human body, is also the predominant bacterial species responsible for community-acquired (CA) and hospital-acquired (HA) infections at all ages in human [1]. Human pathogenic *E. coli* strains are classified into two large groups, strains responsible for intestinal infections and those causing extra-intestinal diseases (ExPEC) [2, 3].

ExPECs are among the most common Gram-negative bacterial pathogens affecting Human with diverse infections, including urinary tract infections (UTI), bacteremia, meningitis, nosocomial respiratory infections, peritonitis, prostatitis, skin and soft tissue infections [4–6].

In addition, multidrug resistant (MDR) ExPECs are now common both in community-acquired and hospital-acquired infections, including resistance to beta-lactams (β-lactams), which are the commonly used antibiotics in human and animal health. The β-lactams resistance is mediated by production of extended-spectrum beta-lactamases [7–9]. These enzymes hydrolyze penicillins, cephalosporins (first, second, third, and fourth generation), and monobactams, but were generally inactive against cephamycins and carbapenems. ESBLs are generally inhibited by beta-lactamase inhibitors (BLI) [10, 11]. A worrying fact is that mobile genetic elements that harbor ESBL genes also carry others genes conferring resistance to quinolones, aminoglycosides and even carbapenems [12–15].

ESBL-producing ExPECs infections are responsible of extended hospital stays, accompanying high cost and mortality and morbidity [2]. Hence, the importance to establish innovative diagnostic toolkits and performant surveillance system for early detection and monitoring of ExPECs cases, especially in developing countries. In this study, we investigated the antibiotic resistance profile and the ESBL genes carried by ESBL-producting ExPEC isolated at the laboratory of bacteriology laboratory, Aristide le Dantec University Teaching Hospital (HALD) in Dakar, Senegal. Additionally, we compared community-acquired (CA) to hospital-acquired (HA), and uropathogenic *E. coli* (UPEC) to no-uropathogenic ExPEC isolates (No-UPEC).

## Materials and methods

### Bacterial isolates

This is a retrospective study and all ExPEC isolates analyzed in this study were collected between January 1^st^, 2018 and December 31^th^, 2020 at the Hospital Laboratory of HALD during routine activities and stored at -80°C. Seventy-eight no-duplicate strains were randomly selected from the Laboratory. Strains were isolated from urine (UPEC, *n* = 51), pus, sputum, bronchial fluid and vaginal samples (no-uropathogenic ExPEC, *n* = 27). Of the 78 strains, 49 and 29 were CA and HA respectively. Culture and Isolation were done based on gold standard microbiological tests and identification by using Api 20E for *Enterobacteriaceae* (bioMérieux France).

### Antibiotic susceptibility testing

Antibiotic susceptibility testing was performed using the Kirby-Bauer disc diffusion method and results were interpreted according to the committee of the French society of microbiology (CA-SFM, 2020) recommendations. Briefly, bacterial suspensions were prepared at 0.5 Mc Farland and inoculated onto Mueller-Hinton agar for an overnight incubation at 37 °C. These following antibiotic disks were tested: ampicillin (AMP, 10 μg), ticarcillin (TIC, 75 μg), amoxicillin-clavulanic acid (AMC, 20/10 μg), cefalotin (CEF, 30 μg), cefoxitin (FOX, 30 μg), cefotaxime (CTA, 30 μg), ceftazidime (CAZ, 30 μg), cefepime (CEP, 30 μg), aztreonam (AZT, 30 μg), imipenem (IMP, 10 μg), ertapenem (ERT, 10μg), Nalidixic acid (NAL, 30 μg), ciprofloxacin (CIP, 5 μg), gentamicin (GEN, 10 μg), amikacin (AMI, 30 μg), fosfomycin (FOS, 50 μg), tetracycline (TET, 30 μg) and sulfamethoxazole-trimethoprim (TMS, 1.25 μg / 23.75 μg). The *E. coli* ATCC 25922 was used for quality control. ESBL production was appreciated by double-disk synergy test with disks of amoxicillin-clavulanic acid surrounded at a radius of 30 mm by cefepime, ceftriaxone, ceftazidime and aztreonam.

### DNA extraction

Bacterial DNA extraction was performed mechanically by thermal choc. Briefly, a well-separated bacterial colony was dispersed in a tube contained 1ml of sterile distilled water, vortexed, boiled for 15 minutes at 100°C and centrifuged at 13,200 rpm for 10 min. The supernatant was carefully recovered, aliquoted and stored at -20°C until used. To confirm results, extraction was done by Qiagen kit (DNeasy Blood & Tissue Kit (50) Cat. No. / ID: 69504).

### ESBL genes amplification

A simplex end-point PCR was performed (on Thermocycler 2720, Applied Biosystems, Lincoln Centre Drive, Foster City, California 94404, USA) to detect ESBL genes. Specific primer pairs (Table 1) were used to amplify ESBL genes (*bla*_CTX-M_, *bla*_CTX-M-1_, *bla*_CTX-M-9_, *bla*_CTX-M-15_, *bla*_CTX-M-25_, *bla*_OXA-1_, *bla*_TEM_, *bla*_SHV_). Each reaction included positive and negative controls. PCRs were carried out in 20 μl reaction volume (2.5 μl DNA + 17.5 Master Mix FIREPol^®^). The amplification program consisted of an initial denaturation at 95°C for 3 min., 35 PCR cycles (denaturation: 94° C, 30 sec., 72°C, 60 sec.) and a final elongation at 72°C for 7 min. Ten microliters of each amplicon were separated on 2% agarose gel in 1X TAE buffer for 35 min at 135 volts and the amplified fragment detected using a GelDoc imager (BioRad).

**Table 1.**
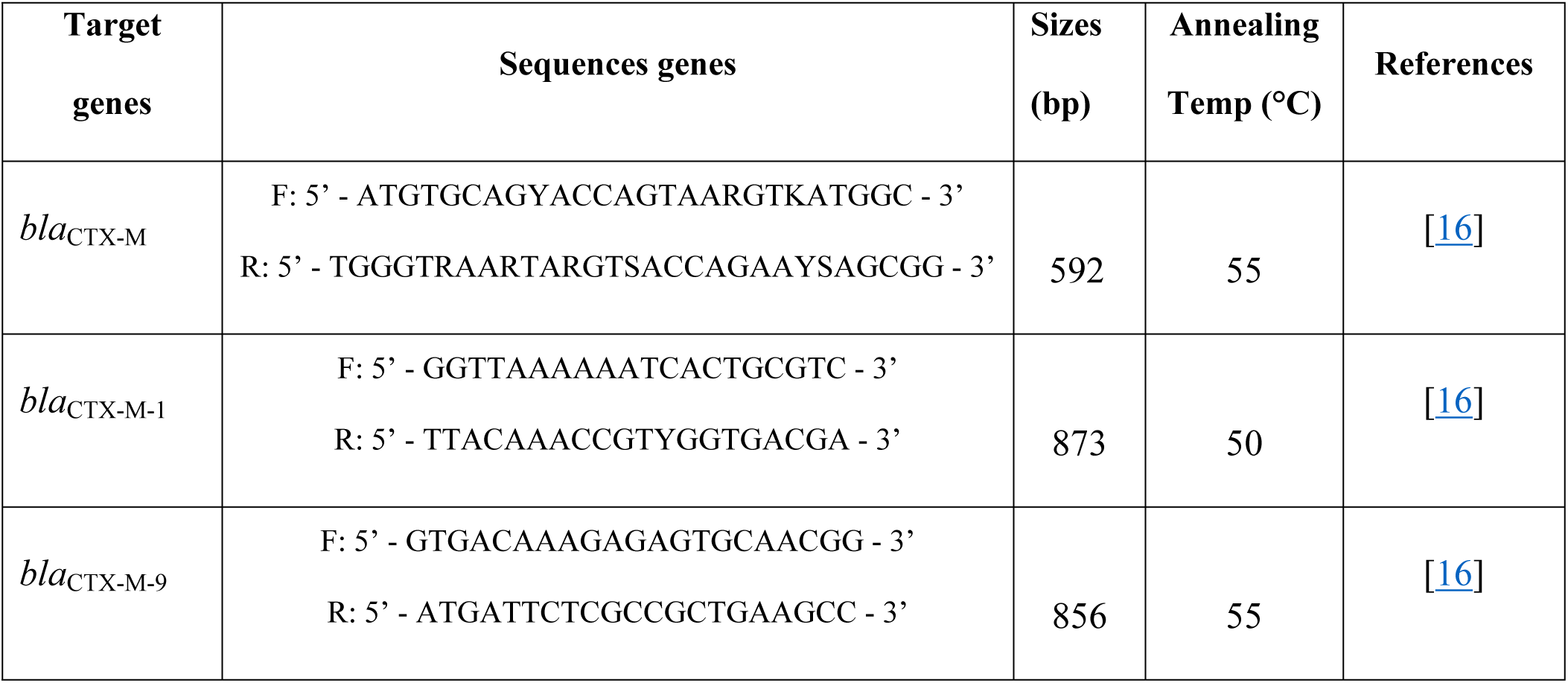
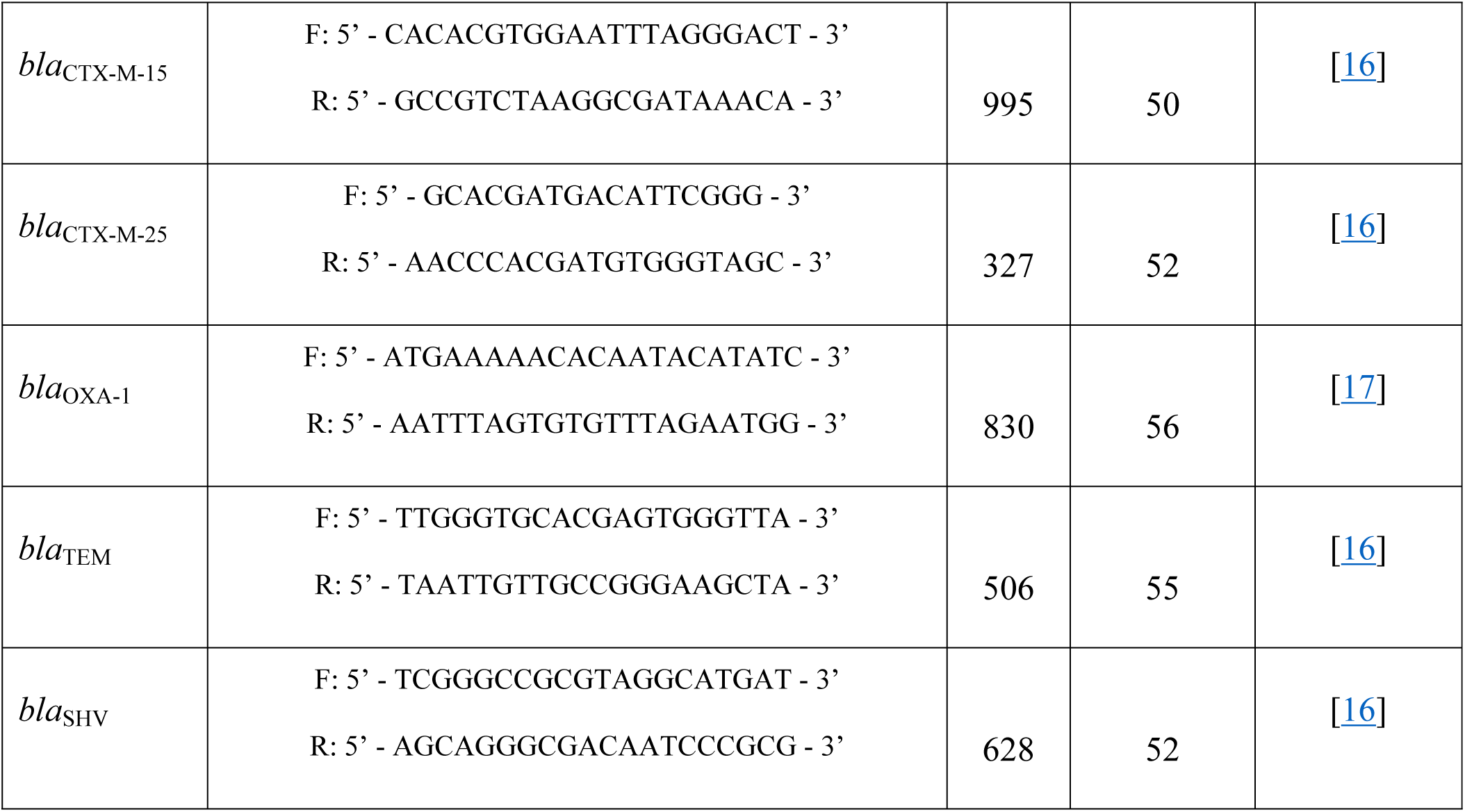
Oligonucleotide primers sequence used for PCR to detect ESBL genes.

### Statistical analysis

Statistical analysis and multiple correspondence analysis and data analysis methods were performed with R software. The statistic test used is the Chi-square at 5% risk threshold. p-values are obtained from the proportion comparison test and the level of significance for all statistical tests was set at p < 0.05.

## Results

### Antibiotic susceptibility testing

All the 78 ExPEC isolates were MDR (resistance to at least one drug from at least three classes of antibiotics), and were resistant to ampicillin, ticarcillin, amoxicillin/clavulanic acid combination, cefalotin, cefotaxime, ceftazidime, cefepime and aztreonam (Table 2). Besides, resistance to ciprofloxacin (93.6%, *n* = 73), tetracycline (91%, *n* = 71) and sulfamethoxazole-trimethoprim (91%, *n* = 71) was high, while less frequent for aminoglycosides (gentamicin, 60.3%, *n* = 47; amikacin, 42.3%, *n* = 33). In contrast, only 3.8% (*n* = 3) and 1.3% (*n* = 1) of the isolates were resistant to ertapenem and fosfomycin respectively, while all were sensitive to imipenem (Table 2). Comparison of resistance profiles between CA and HA, and between UPEC and no-UPEC strain did not show any significant difference, except for ciprofloxacin (Table 2).

**Table 2.** Antibiotics resistance rate of total strains, CA and HA strains, UPEC and no-UPEC strains.

| Antibiotics |  | Total strains | Pathogenicity |  |  | Origin |  |  |
| --- | --- | --- | --- | --- | --- | --- | --- | --- |
| Class | Drug | N (%) | UPEC<br>N (%) | No-UPEC<br>N (%) | p | CA<br>N (%) | HA<br>N (%) | p |
| Beta-lactams | AMP | 78 (100) | 51 (100) | 27 (100) | 1 | 49 (100) | 29 (100) | 1 |
|  | TIC | 78 (100) | 51 (100) | 27 (100) | 1 | 49 (100) | 29 (100) | 1 |
|  | AMC | 78 (100) | 51 (100) | 27 (100) | 1 | 49 (100) | 29 (100) | 1 |
|  | CEF | 78 (100) | 51 (100) | 27 (100) | 1 | 49 (100) | 29 (100) | 1 |
|  | FOX | 5 (6.4) | 3 (5.9) | 2 (7.4) | 0.06 | 4 (8.2) | 1 (3.5) | 0.06 |
|  | CTA | 78 (100) | 51 (100) | 27 (100) | 1 | 49 (100) | 29 (100) | 1 |
|  | CAZ | 78 (100) | 51 (100) | 27 (100) | 1 | 49 (100) | 29 (100) | 1 |
|  | CEP | 78 (100) | 51 (100) | 27 (100) | 1 | 49 (100) | 29 (100) | 1 |
|  | AZT | 70 (89.7) | 48 (91.4) | 22 (81.5) | 0.08 | 46 (93.9) | 24 (82.8) | 0.12 |
|  | IMP | 0 | 0 | 0 | - | 0 | 0 | - |
|  | ERT | 3 (3.8) | 3 (5.9) | 0 | 0.01* | 2 (4.1) | 1 (3.5) | 0.04* |
| Quinolones and<br>Fluoroquinolones | NAL | 76 (97.4) | 51 (100) | 25 (92.6) | 0.97 | 49 (100) | 27 (93.1) | 0.97 |
|  | CIP | 73 (93.6) | 50 (98) | 23 (85.2) | 0.03* | 48 (98) | 25 (86.2) | 0.04* |
| Aminoglycosides | GEN | 47 (60.3) | 33 (64.7) | 14 (51.9) | 0.27 | 29 (59.2) | 16 (62.1) | 0.72 |
|  | AMI | 33 (42.3) | 24 (47.1) | 9 (33.3) | 0.24 | 19 (38.8) | 14 (48.3) | 0.41 |
| Phosphonic acid | FOS | 1 (1.3) | 1 (2) | 0 | 0.01* | 0 | 1 (3.6) | 0.01* |
| Cyclines | TET | 71 (91) | 45 (88.2) | 26 (96.3) | 0.24 | 44 (89.8) | 27 (93.1) | 0.62 |
| Antifolates | TMS | 71 (91) | 45 (88.2) | 26 (96.3) | 0.24 | 45 (91.8) | 26 (89.7) | 0.74 |

UPEC, Uropathogenic *E. coli*; CA, Community-acquired; HA, Hospital-acquired; AMP, ampicillin; TIC, ticarcillin; AMC, Amoxicillin-clavulanic acid; CEF, cefalotin; FOX, cefoxitin; CTA, cefotaxim; CAZ, ceftazidime; CEP, cefepime; AZT, aztreonam; IMP, imipenem; ERT, Ertapenem; NAL, nalidixic acid; CIP, ciprofloxacin; GEN, gentamicin; AMI, amikacin; FOS, fosfomycin; TET, tetracycline; TMS, sulphamethoxazole-trimethoprim; \*, significant p-value ( $p < 0.05$ ).

### Presence of ESBL genes

All 78 strains carried at least one ESBL gene. *bla*_CTX-M_ group was the most prevalent (77/78; 98.7%), followed by *bla*_OXA-1_ (61/78; 78.2%), *bla*_TEM_ (35/78; 44.9%) and *bla*_SHV_ (3/78; 3.8%) (Table 3) and (Fig 1). 51/51 (100%) of UPEC strains and 29/29 (100%) of hospital -acquired strains carried the *bla*_CTX-M_ gene and none of ‘‘no-uropathogenic ExPEC‘‘ strains carried a *bla*_SHV_ gene (Table 3) and (Fig 2). (9/78; 11.5%) carried only *bla*_CTX-M_ or *bla*_OXA-1_ and (69/78; 88.5%) carried several types of ESBL gene. Indeed, (2/78; 2.6%) carried *bla*_CTX-M_ + *bla*_OXA-1_ + *bla*_TEM_ + *bla*_SHV_; (23/78; 29.4%) carried *bla*_CTX-M_ + *bla*_OXA-1_ + *bla*_TEM_; (1/78; 1.3%) carried *bla*_CTXM_ + *bla*_OXA-1_ + *bla*_SHV_; (32/78; 41%) carried *bla*_CTX-M_ + *bla*_OXA-1_ and (11/78; 14.1%) of strains carried “*bla*_CTX-M_ + *bla*_TEM_” (Table 3). None of strains carried *bla*_TEM_ or *bla*_SHV_ gene alone (Table 4).

**Table 3.** Prevalence of ESBL genes in total strains, CA and HA strains, UPEC and non-UPEC strains.

| ESBL |  | Total strains | Pathogenicity |  |  | Origin |  |  |
| --- | --- | --- | --- | --- | --- | --- | --- | --- |
| Family | Genes | N (%) | UPEC<br>N (%) | No-UPEC<br>N (%) | p | CA<br>N (%) | HA<br>N (%) | p |
| Cefotaximase-Munich | *bla*<sub>CTX-M</sub> | 77 (98.7) | 51 (100) | 26 (96.3) | 0.98 | 48 (98) | 29 (100) | 0.99 |
|  | *bla*<sub>CTX-M-1</sub> | 70 (89.7) | 48 (94.1) | 22 (81.5) | 0.08 | 44 (89.8) | 26 (89.7) | 0.98 |
|  | *bla*<sub>CTX-M-9</sub> | 9 (11.5) | 5 (9.8) | 4 (14.8) | 0.25 | 5 (10.2) | 4 (13.8) | 0.45 |
|  | *bla*<sub>CTX-M-15</sub> | 70 (89.7) | 48 (94.1) | 22 (81.5) | 0.08 | 44 (89.8) | 26 (89.7) | 0.98 |
|  | *bla*<sub>CTX-M-25</sub> | 0 | 0 | 0 | - | 0 | 0 | - |
| Oxacillinase | *bla*<sub>OXA-1</sub> | 61 (78.2) | 42 (82.4) | 19 (70.4) | 0.22 | 38 (77.6) | 23 (79.3) | 0.86 |
| Temoneira | *bla*<sub>TEM</sub> | 35 (44.9) | 23 (45.1) | 12 (44.4) | 0.95 | 19 (38.8) | 16 (55.2) | 0.16 |
| Sulfhydryl variable | *bla*<sub>SHV</sub> | 3 (3.8) | 3 (5.9) | 0 | 0.01* | 2 (4.1) | 1 (3.5) | 0.04* |
| UPEC, Uropathogenic *E. coli*; CA, community-acquired; HA, hospital-acquired; %, percentage; N, number of isolates, *, significant p-value (< 0.05). |  |  |  |  |  |  |  |  |

**Table 4.** Prevalence of ESBL genes combinations in total strains, CA and HA strains, UPEC and non-UPEC strains.

| Combination of ESBL genes | Total strains | Pathogenicity |  |  | Origin |  |  |
| --- | --- | --- | --- | --- | --- | --- | --- |
|  | N (%) | UPEC<br>N (%) | No-UPEC<br>N (%) | p | CA<br>N (%) | HA<br>N (%) | p |
| *bla*<sub>CTX-M-1</sub> + *bla*<sub>CTX-M-15</sub> + *bla*<sub>CTX-M-9</sub><br>+ *bla*<sub>OXA-1</sub> + *bla*<sub>TEM</sub> | 2 (2.6) | 2 (3.9) | 0 | 0.01* | 1 (2) | 1 (3.4) | 0.1 |
| *bla*<sub>CTX-M-1</sub> + *bla*<sub>CTX-M-15</sub> + *bla*<sub>OXA-1</sub> +<br>*bla*<sub>TEM</sub> + *bla*<sub>SHV</sub> | 2 (2.6) | 2 (3.9) | 0 | 0.01* | 1 (2) | 1 (3.4) | 0.1 |
| *bla*<sub>CTX-M-1</sub> + *bla*<sub>CTX-M-15</sub> + *bla*<sub>OXA-1</sub> +<br>*bla*<sub>TEM</sub> | 21 (26.9) | 14 (27.4) | 7 (33.3) | 0.68 | 10 (20.4) | 11 (37.9) | 0.12 |
| *bla*<sub>CTX-M-1</sub> + *bla*<sub>CTX-M-15</sub> + *bla*<sub>OXA-1</sub> +<br>*bla*<sub>SHV</sub> | 1 (1.3) | 1 (2) | 0 | 0.01* | 1 (2) | 0 | 0.01* |
| *bla*<sub>CTX-M-1</sub> + *bla*<sub>CTX-M-15</sub> + *bla*<sub>OXA-1</sub> | 32 (41) | 21 (41.2) | 11 (40.7) | 0.98 | 23 (46.9) | 9 (31) | 0.15 |
| *bla*<sub>CTX-M-1</sub> + *bla*<sub>OXA-1</sub> + *bla*<sub>TEM</sub> | 1 (1.3) | 1 (2) | 0 | 0.01* | 1 (2) | 0 | 0.01* |
| *bla*<sub>CTX-M-1</sub> + *bla*<sub>CTX-M-15</sub> + *bla*<sub>TEM</sub> | 7 (9) | 4 (7.8) | 3 (11.1) | 0.21 | 5 (10.2) | 2 (6.9) | 0.23 |
| *bla*<sub>CTX-M-1</sub> + *bla*<sub>CTX-M-15</sub> | 3 (3.8) | 2 (3.9) | 1 (3.7) | 0.97 | 2 (4.1) | 1 (3.4) | 0.04 |
| *bla*<sub>CTX-M-15</sub> + *bla*<sub>CTX-M-9</sub> | 2 (2.6) | 2 (3.9) | 0 | 0.01* | 1 (2) | 1 (3.4) | 0.1 |
| *bla*<sub>CTX-M-1</sub> + *bla*<sub>OXA-1</sub> | 1 (1.3) | 1 (2) | 0 | 0.01* | 0 | 1 (3.4) | 0.01* |
| *bla*<sub>CTX-M-9</sub> + *bla*<sub>TEM</sub> | 2 (2.6) | 0 | 2 (7.4) | 0.01* | 1 (2) | 1 (3.4) | 0.1 |
| UPEC, Uropathogenic *E. coli*; CA, community-acquired; HA, hospital-acquired %, percentage; N, number of isolates;<br>*, significant p-value (< 0.05). |  |  |  |  |  |  |  |

**Fig. 1.**
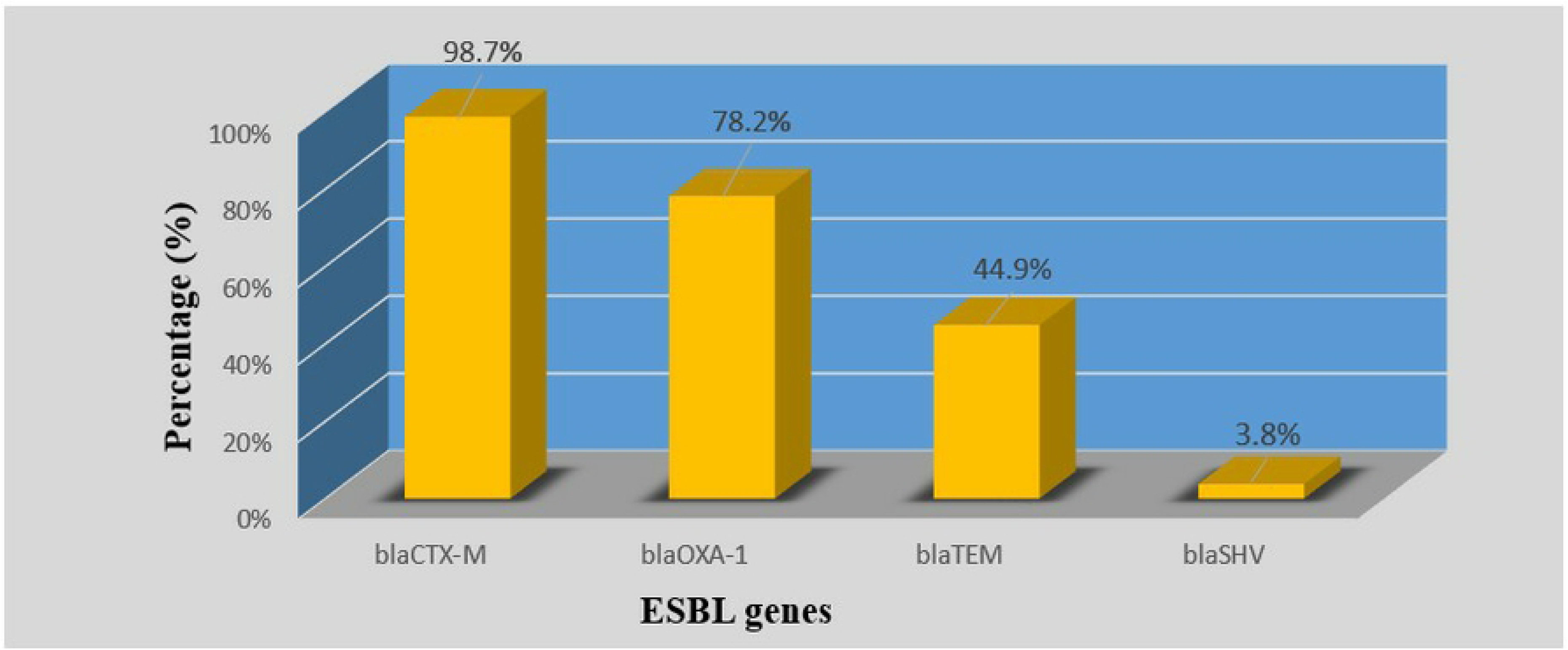
Prevalence of ESBL genes in total strains.

**Fig. 2.**
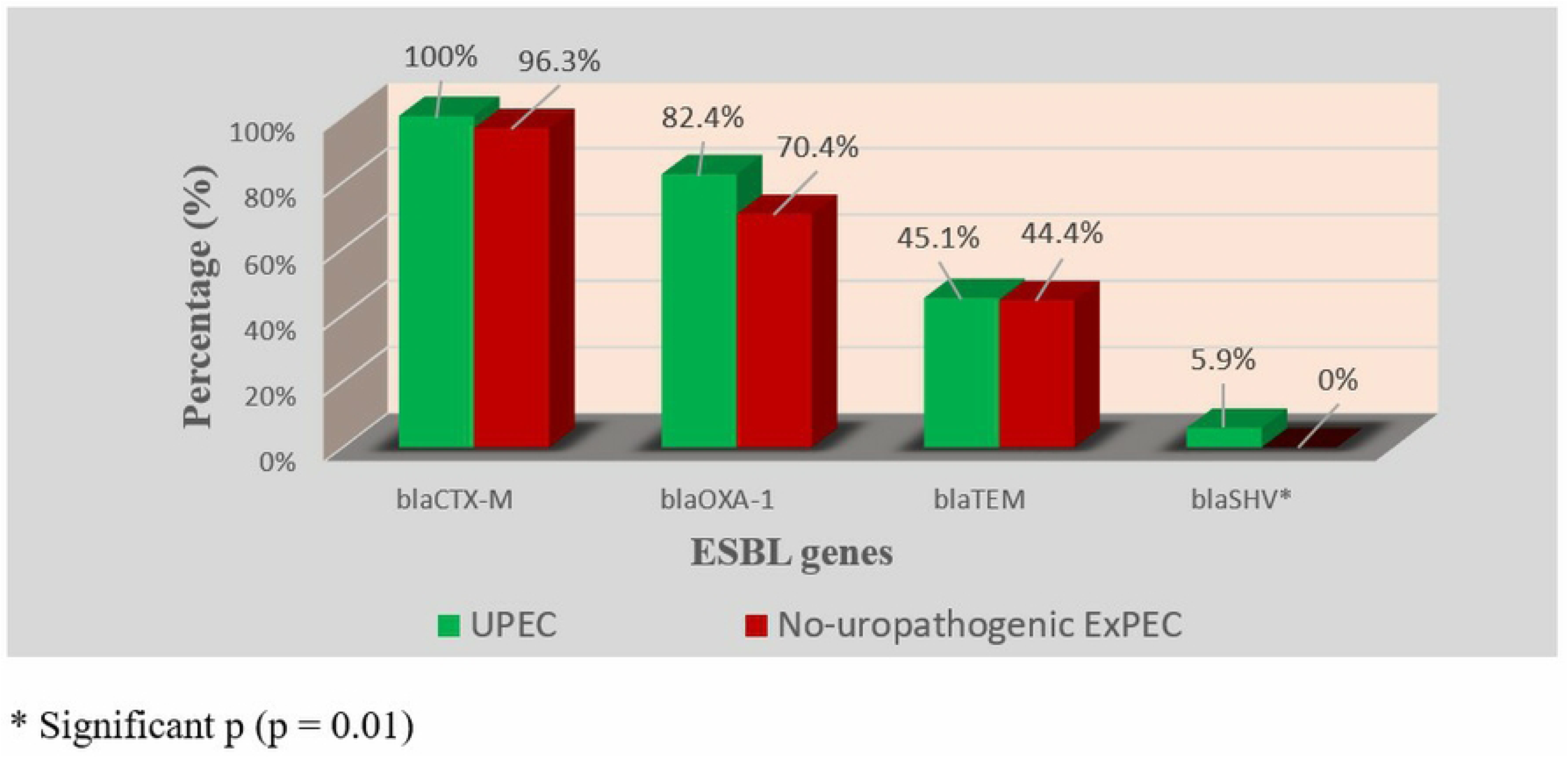
Prevalence of ESBL genes in UPEC and no-uropathogenic ExPEC.

In 4 *bla*_CTX-M_ group, *bla*_CTX-M-1_ (70/77; 90.9%) with *bla*_CTX-M-15_ (70/77; 90.9%) was the most prevalent followed by *bla*_CTX-M-9_ (9/77; 11.7%). *bla*_CTX-M-25_ was not detected in any of the 77 strains (Table 4). Among strains which carried the *bla*_CTX-M_ type, 89.6% carried 2 variants of *bla*_CTX-M_ while 7.8% carried only one variant and 2.6% carried 3 *bla*_CTX-M_ variants (Table 4). No significant difference was found by comparing the prevalence of ESBL genes in hospital-acquired and community-acquired strains on the one hand and UPEC and no-uropathogenic ExPEC strains on the other hand (Figs 2-5).

**Fig. 3.**
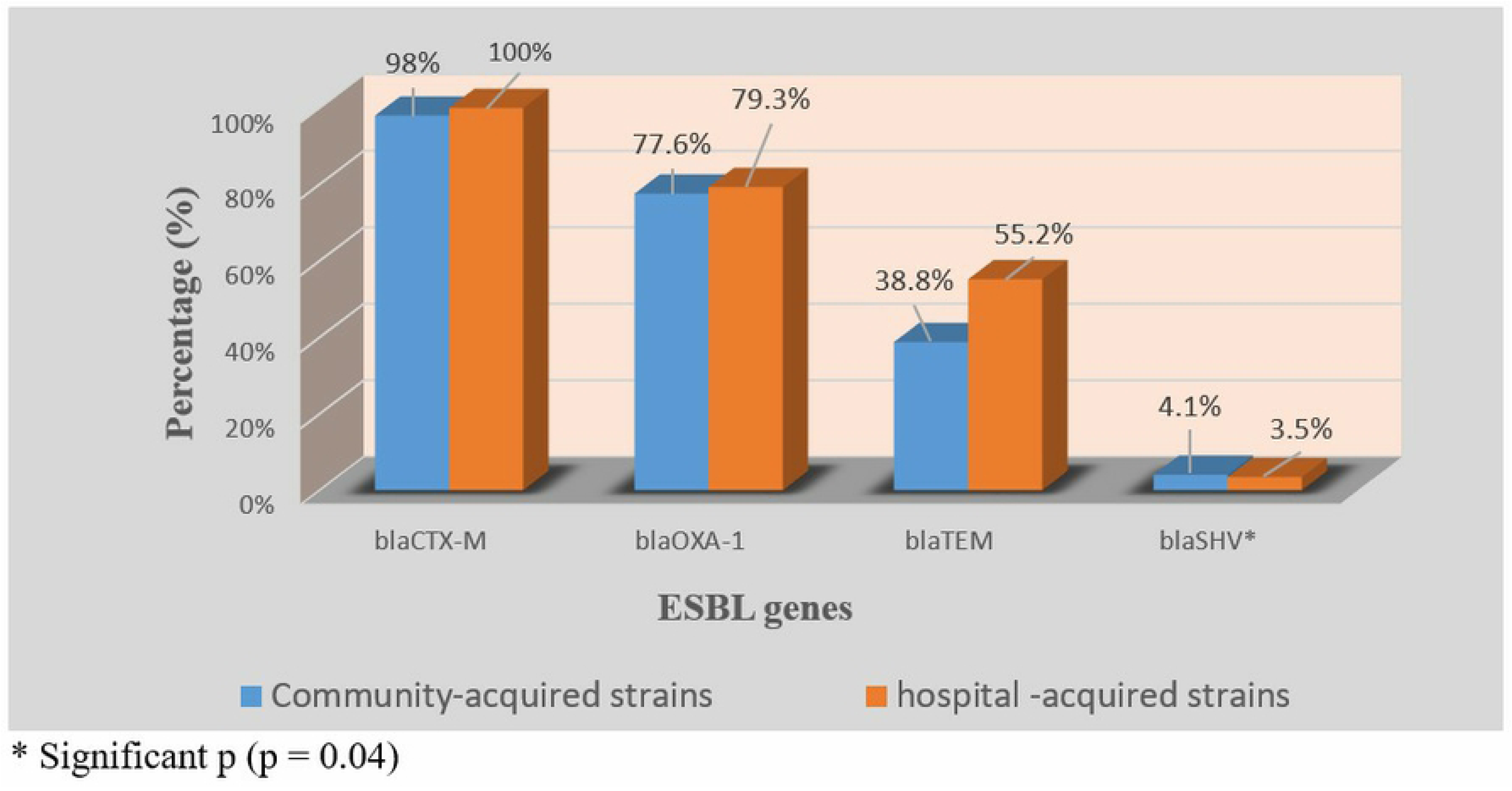
Prevalence of ESBL genes in community-acquired and hospital-acquired strains.

**Fig. 4.**
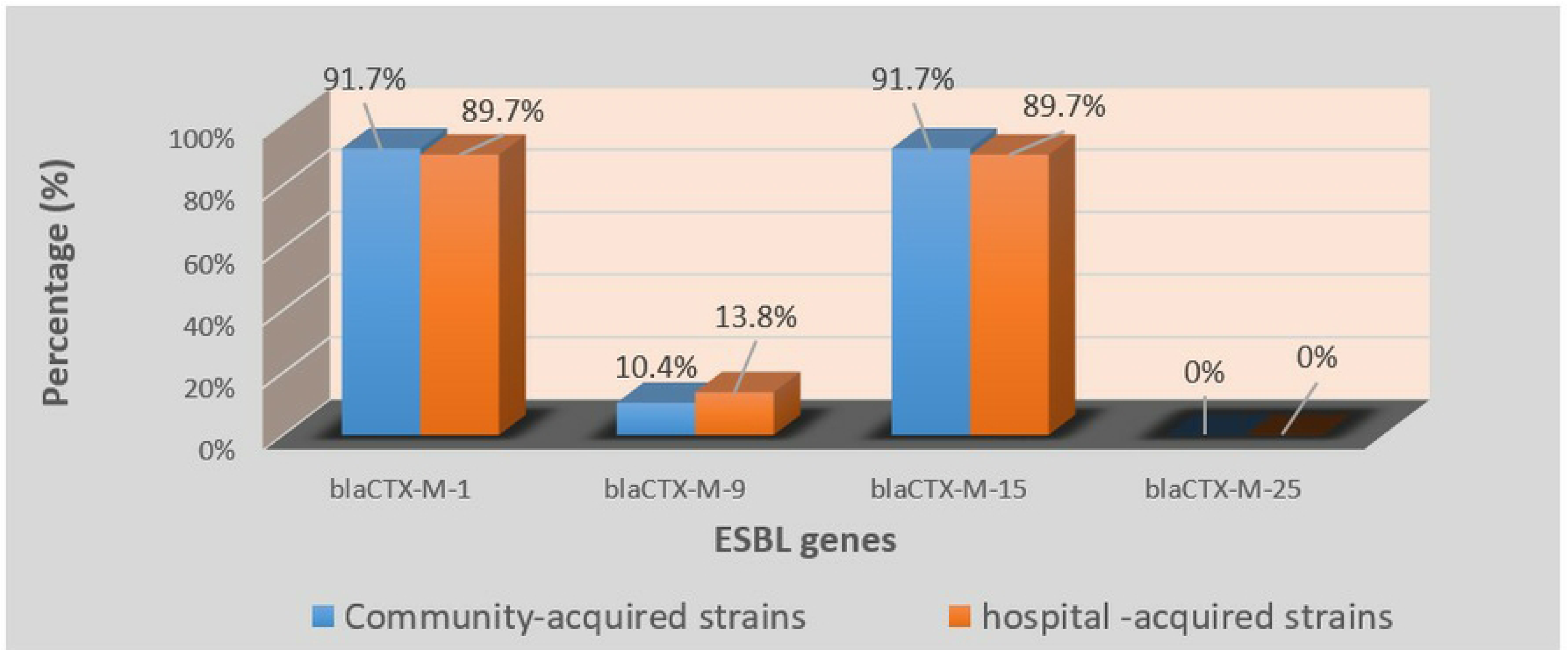
Prevalence of bla_CTX-M_ variants in community and hospital-acquired strains.

**Fig. 5.**
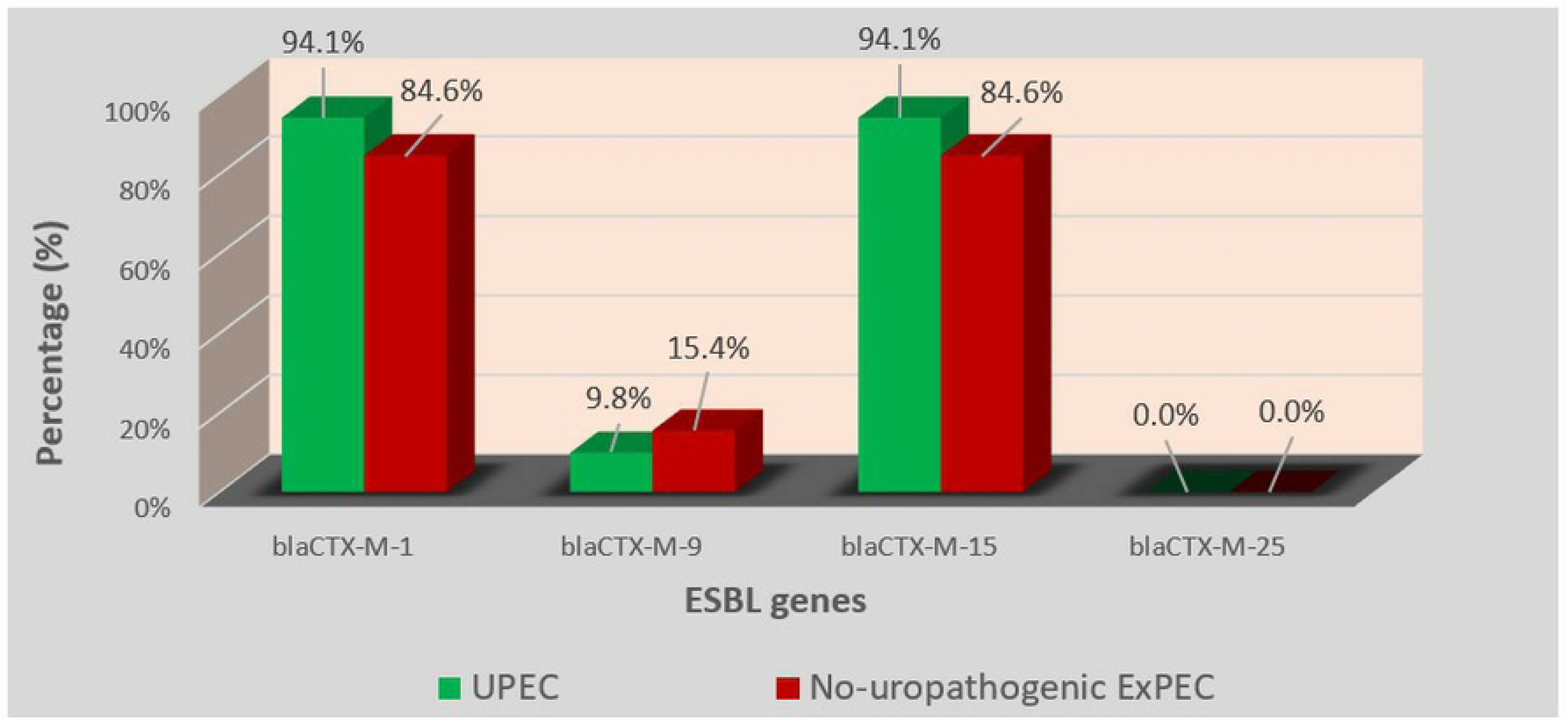
Prevalence of bla_CTX-M_ variants genes in UPEC and no-uropathogenic ExPEC.

## Discussion

Potentially pathogenic *Escherichia coli* which produce extended spectrum betalactamases (ESBL) are frequently isolated from urinary tract infections [6, 18] can be resistant to many molecules of this class. The most worrying thing is that these strains, which spread rapidly both in hospitals and in the community, are often resistant to many other antibiotics such as those of the aminoglycosides and quinolones classes, thus making the treatment failure of these infections.

The rate of resistance to multiple antibiotics among ESBL-producing isolates is usually common due to carrying multi-resistant genes and plasmids [12–14].

Nowadays, ESBL type CTX-M are the most widespread in the world, unlike ESBL TEM and SHV which are becoming less prevalent [19, 20]. Our study also confirmed this trend with 98.7% of strains positive for *bla*_CTX-M_, followed by *bla*_OXA-1_ (78.2%), *bla*_TEM_ (44.9%) and *bla*_SHV_ (3.8%). Several studies carried out in Togo [21], in Saudi Arabia [22], and in Mozambique [23] reported high prevalence rates of *bla*_CTX-M_ in ESBL ExPEC strains 100%; 93,94% and 77%, respectively. Moreover, other authors [24, 25] had already pointed out that currently, *bla*_OXA-1_ was the second most prevalent ESBL gene type in the world behind *bla*_CTX-M_. In disagreement to these studies, [23] rather mentioned 52% of prevalence for *bla*_SHV_ and 1% for *bla*_TEM_ in 2021 in Mozambique. [26] mentioned 3.12% of prevalence rate for *bla*_SHV_ in 2019 in Senegal. ESBLs SHV-gene type therefore seem to be rare in *E. coli* strains circulating in Senegal.

Interestingly, 55.1% of the strains harbored 2 ESBL gene types while 30.8% of the strains carried 3 and 2.6% carried all the 4 ESBL genotypes. While 33.33% and 12.12% of strains carrying 2 and 3 ESBL gene types, respectively were reported from Riyadh, Saudi Arabia [22]. The very high proportion of strains (88.5%) combining several ESBL gene types seems to be one of the major causes of the 100% resistance to ampicillin, ticarcillin, (clavulanic acid + amoxicillin), cefalotin, cefotaxime, ceftazidime, cefepime and aztreonam. None of the strains carried only *bla*_TEM_ or *bla*_SHV_ gene.

Globally, *bla*_CTX-M-15_ had long been cited as the most prevalent variant of *bla*_CTX-M_ in *E. coli* [27–29]. The high prevalence rate of *bla*_CTX-M-15_ (90.9%) observed among *bla*_CTX-M_ positive strains in our study corroborates these earlier studies. An interesting fact in our study was that *bla*_CTX-M-1_ was as prevalent as *bla*_CTX-M-15_ with respectively 90.9% and 85.7% rates of the strains concomitantly carried *bla*_CTX-M-15_ and *bla*_CTX-M-1_. These data suggest that *bla*_CTX-M-15_ is not the only major variant of *bla*_CTX-M_ circulating in Senegal. The low prevalence rates of *bla*_CTX-M-9_ (11.7%) and *bla*_CTX-M-25_ (0%) follow trends observed in other parts of the world [27, 30].

No significant difference was noted when comparing the prevalence of ESBL genes from community-acquired and hospital-acquired strains. This seems to imply either a port of ESBL ExPEC in community or that the community strains are the same ones encountered in a hospital environment. We did not notice any significant difference in the prevalence of ESBL genes between UPEC strains and non-uropathogenic ExPEC strains. It seems that in Senegal, non-uropathogenic ExPEC are as resistant as UPEC strains. Future studies could confirm this.

The high prevalence of *bla*_CTX-M_ genes suggests the involvement of mobile genetic elements (plasmids, integrons and transposons) [20, 31] in the spread of antibiotic resistance in Dakar, as reported in many studies leading increasing resistance to fluoroquinolones, aminoglycosides and even carbapenems antibiotics [12–14]. This suggests the importance to study and monitor the mobile genetic elements from strains isolated in healthy carriers, environment, and hospital settings in order to initiate others actions that can help fighting against antibiotic resistance.

## Conclusion

All the 78 ExPEC strains tested in this study were MDR patterns, and resistant to almost all antibiotics families, except fosfomycin and carbapenems. Based on our results, we recommend avoiding monotherapy and prohibiting fluoroquinolones, C3G and C4G as empiric treatment of UTIs in Senegal. *bla*_CTX-M_ (*bla*_CTX-M1_, *bla*_CTX-M15_) and *bla*_OXA-1_ seem to be the major ESBL genes circulating in Senegal. No significant difference was noted when comparing the prevalence of ESBL genes between hospital-acquired and community-acquired strains; As well as by comparing UPEC and ExPEC strains isolated from other types of samples. The high resistance to antimicrobials observed, underscore the relevance to implement an epidemiological antimicrobial resistance (AMR) surveillance system to improve the management of treatment protocols in patients infected with MDR bacteria.

## Acknowledgements

The authors thank Abdoul Aziz Wane, Amadou Mactar Gueye, Ousmane Sow and El Hadji Aly Niang for their technical assistance.

## Notes

### Competing Interest Statement

The authors have declared no competing interest.

